# CHD8 adulthood microglial knockout induces behavioral, morphological, and transcriptional changes in a sex-dependent manner

**DOI:** 10.1101/2024.06.16.599211

**Authors:** Orly Weissberg, Dmitrity Getselter, Evan Elliott

## Abstract

Mutations in CHD8 (chromodomain-helicase-DNA binding protein 8) are highly associated with autism spectrum disorders. It has been well established that CHD8 has a prominent role in the development of neurons. However, there is little knowledge of its specific roles in microglia, and its possible roles in cellular functions after development, i.e. adulthood. In addition, while microglial dysfunction has been characterized in autism, the roles of autism-associated genes in microglial function have not been well characterized. Using conditional transgenic mouse models, we determined that adulthood deletion of CHD8 in microglia induces robust changes in behavior, including anxiety, social deficits, and depression-like behavior, in association with changes in microglial activation and robust microglial gene expression changes, including expression of cytokines. Of great interest, many of these changes were seen specifically in male deletion mice, and not female deletion mice. In contrast, adulthood neuron knockout had more subtle effects on behavior, mainly on depression-like behavior, and induced subtle changes in gene transcription related to the Wnt/Beta-Catenin pathway. These changes were also only present in male neuron knockout mice. In summary, CHD8 is particularly important for microglial function in adulthood and has cellular effects that are specific to males.

## Introduction

Autism spectrum disorder is a group of neurodevelopmental disorders that begin in early childhood. Behavior patterns and etiologies of autism are highly heterogeneous^1,2^ and prevalence is four times higher among boys than girls. Clinical diagnosis is characterized by impaired social communication and language development, as well as repetitive and stereotypical behaviors, and prevalence has risen during the last decades to 1 out of 44 eight-year-old children in the USA^3^ and about 1% of the worldwide population4. The etiology of autism is complex, involving both genetics and environmental factors^2,5–7^. Co-morbidities of autism include anxiety, depression, attention deficit hyperactivity disorder (ADHD), speech and language delay, sleep disorders, gastrointestinal problems, and impaired learning^8–10^.

Chromodomain helicase DNA-binding protein 8 (CHD8) is located on chromosome 14q11.2 in humans and has two isoforms, a large isoform, and a small isoform ^11,12^. CHD8 mutations are among the most common mutations associated with autism and cause approximately 0.5% of all cases. Interestingly, unlike many other autism-related genes, CHD8 mutations have not been found in typical control groups, suggesting that CHD8 has high penetrance^13–17^. In particular, many de novo mutations in CHD8 were found to be part of autism etiology^10,14,18,19^. Individuals carrying CHD8 mutations have unique phenotypes including atypical face, macrocephaly, tall and slender stature, gastrointestinal disturbance, anxiety, and sleep problems^10,20–24^.

CHD8 is a chromatin-binding factor and was found to directly interact with β-catenin and negatively regulate its targeted genes including WNT signaling ^11,25^. In addition, CHD8 was found to regulate the expression of many genes considered to be autism risk factors and involved in human neurodevelopment. According to ChIP-seq analysis, CHD8 directly binds genomic sites in neurodevelopment-related and autism-risk genes^14,26^. Interestingly, mutations in CHD8 show a male bias effect (in a ratio of ∼85:15) but the reason is unclear^16,27–32^

Several CHD8 knockout mouse models have been developed to study the role of CHD8 in brain function and development. However, most studies investigated the effect of CHD8 mutations in males only^10,28,31,33–40^. CHD8 Haploinsufficient developmental models displayed increased total brain volume^28,31,33,35,36^, increased absolute volume of the cerebral cortex, hippocampus and amygdala^35,36^ increase cortical and hippocampal connectivity (rsfMRI)^36^ and sexually dimorphic neuronal firing and synaptic transmission in the hippocampus along with normal brain morphology (MRI)^28^. Chd8 has an important role in the cell proliferation of the embryonic neocortex^40^. However, nearly all these studies determined the roles of CHD8 in neuronal development and did not look into possible roles for CHD8 after developmental time points.

Within the central nervous system, microglia are the primary phagocytotic cells, accounting for 5-20% of all glial cells^41^. They play a critical role in brain development ^42^ and are involved in neurogenesis, axonal migration, synapse formation, and programmed cell death^43,44^. As integral components of the immune system, they play a crucial role in regulating neuroinflammatory responses^41,45,46^. Microglia were found to be involved in many neurodevelopmental disorders including autism^42,47–53^. Initial human studies found evidence for microglial activation in children diagnosed with autism^54^. Furthermore, animal studies have found that environmental factors, such as maternal immune activation during pregnancy, can induce activation of microglia associated with behavioral and neurodevelopmental changes in offspring^55^. Therefore, microglial dysregulation has been proposed as a biological mechanism linking environmental factors to autism. However, few studies have studied the link between genetic factors involved in autism and microglial regulation.

While CHD8 has been well studied at early developmental time points, it not well known if CHD8 continues to have roles in post-developmental time points. Therefore, it is not known if development of tools to induce CHD8 levels after early development could be relevant for treatment in individuals with mutations in CHD8.

This study aimed to shed light on several aspects of CHD8 regulation in the brain, with a primary focus on its role during adulthood and its specific roles in neurons and microglia in adulthood. In addition, we aim to decipher the sex-specific effects of CHD8 in these cellular subtypes.

## Results

### Generation and Characterization of Adulthood-Specific CHD8 Knockout Mice in Microglia and Excitatory Neurons

To generate adulthood-specific knockout models of CHD8 in distinct cell populations, CHD8^flox^ mice were crossbred with Cx3cr1^Cre-ERT^/CHD8^flox^ or CamKIIa^Cre-ERT^ /CHD8^flox^ transgenic lines (Fig. 1a). Inducible microglial CHD8 knockout (imKO mice) were obtained by tamoxifen administration to 8-week-old CHD8^flox^/Cx3cr1^Cre-ERT+^ mice. It is important to note that in order to induce a microglia-specific knockout, tamoxifen treatment must be done in the adulthood (after eight weeks of age) because is Cx3cr1 is expressed at multiple tissues during early development. Tamoxifen-treated CHD8^flox^/Cx3cr1^Cre-ERT-^ mice were used as controls. Four weeks post-treatment, immunohistochemical analysis confirmed the reduction of Chd8 expression specifically in microglial cells in the Prefrontal Cortex (PFC) (Fig. 1b). Concurrently, adult-onset CHD8 knockout in excitatory neurons (inKO mice) was achieved by administering tamoxifen to 8-week-old CHD8^flox^/CamKIIa^Cre-ERT+^ mice and CHD8^flox^/CamKIIa^Cre-ERT-^ controls. Immunohistochemical analysis performed four weeks post-treatment revealed a robust Chd8 depletion distinctively within neurons in the hippocampus (Fig. 1c).

**Figure 1.**
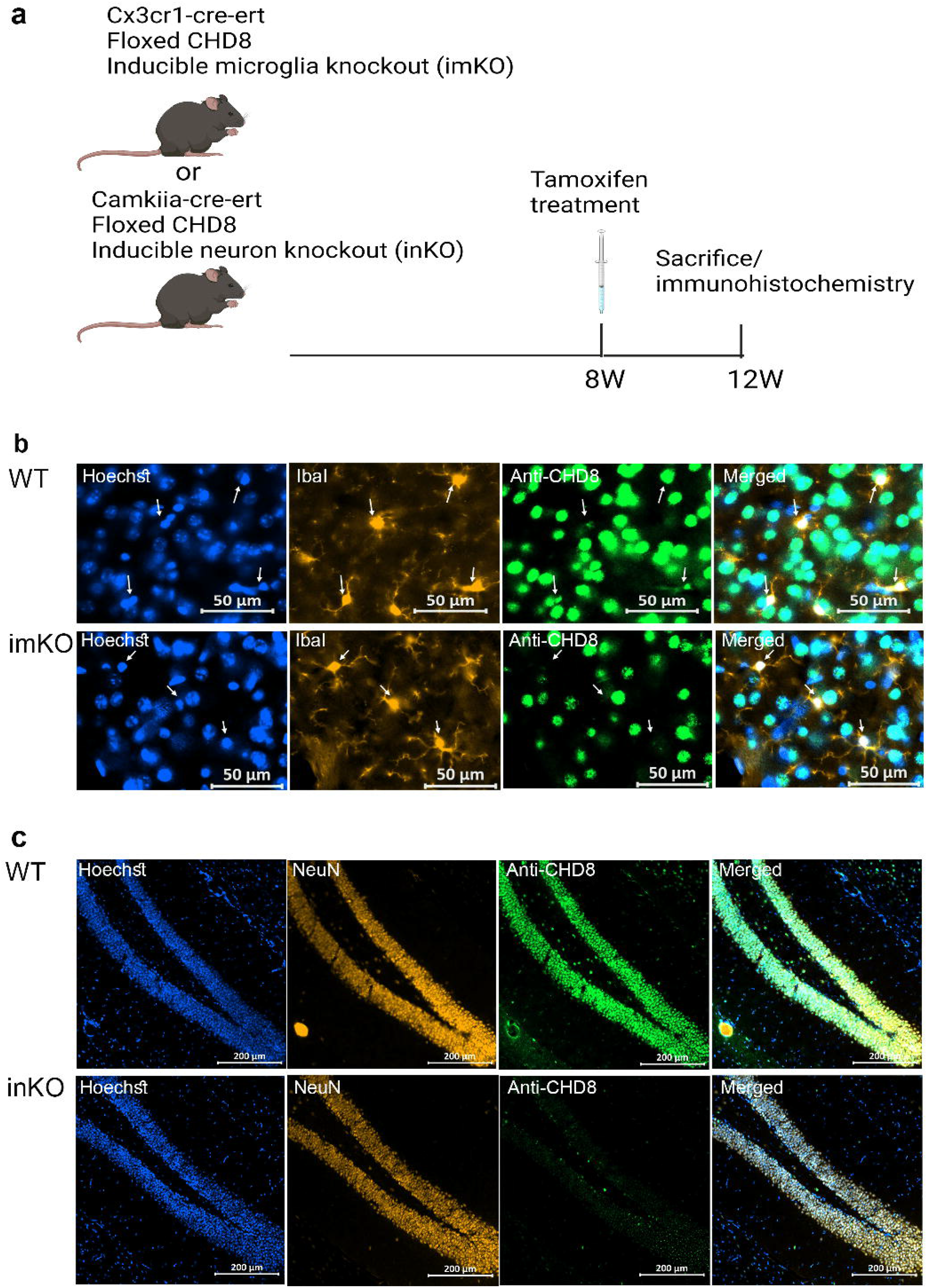
Characterization of CHD8 depletion in animal models. a) Timeline of tamoxifen administration and animal sacrifice of two animal models for immunohistochemistry analysis. b) Immunohistochemistry of prefrontal cortex with antibodies targeting microglia (IbaI positive), and CHD8. Nuclei are stained in blue (Hoechst). c) Immunohistochemistry of hippocampus with antibodies targeting neurons (NeuN), and CHD8. Nuclei are stained in blue (Hoechst).

### Sex-Specific Effect of CHD8 on imKO and inKO Mice

A panel of behavioral tests were performed on the imKO (microglia-specific knockout) in both male and female cohorts. Male imKO mice exhibited reduced time spent in the center of the open field (Fig. 2a) and in the light zone of the dark-light chamber (Fig. 2b) compared to wild type controls. Time spent in the open arms of the elevated plus maze was not significantly different between the two genotypes (Fig. 2c). There were no differences between the genotypes in tests for repetitive behaviors (Fig. 2d,e). However, male imKO mice displayed depression-like behaviors evident by increased immobility in the tail suspension test (Fig. 2f) and forced swimming test compared to wild type controls (Fig. 2g). There were now difference between genotypes in general motor function in the rotarod test (Fig. 2h). In the cue fear conditioning test (Fig. 2i), imKO male mice had reduced freezing responses, indicating possible memory and learning impairments. Moreover, alterations in sociability behaviors were evident, as imKO male mice exhibited a lack of preference for the stranger versus an empty cage (Fig. 2j). Therefore, male imKO mice displayed several behavior phenotypes including an increase in anxiety and depression-like behavior.

**Figure 2.**
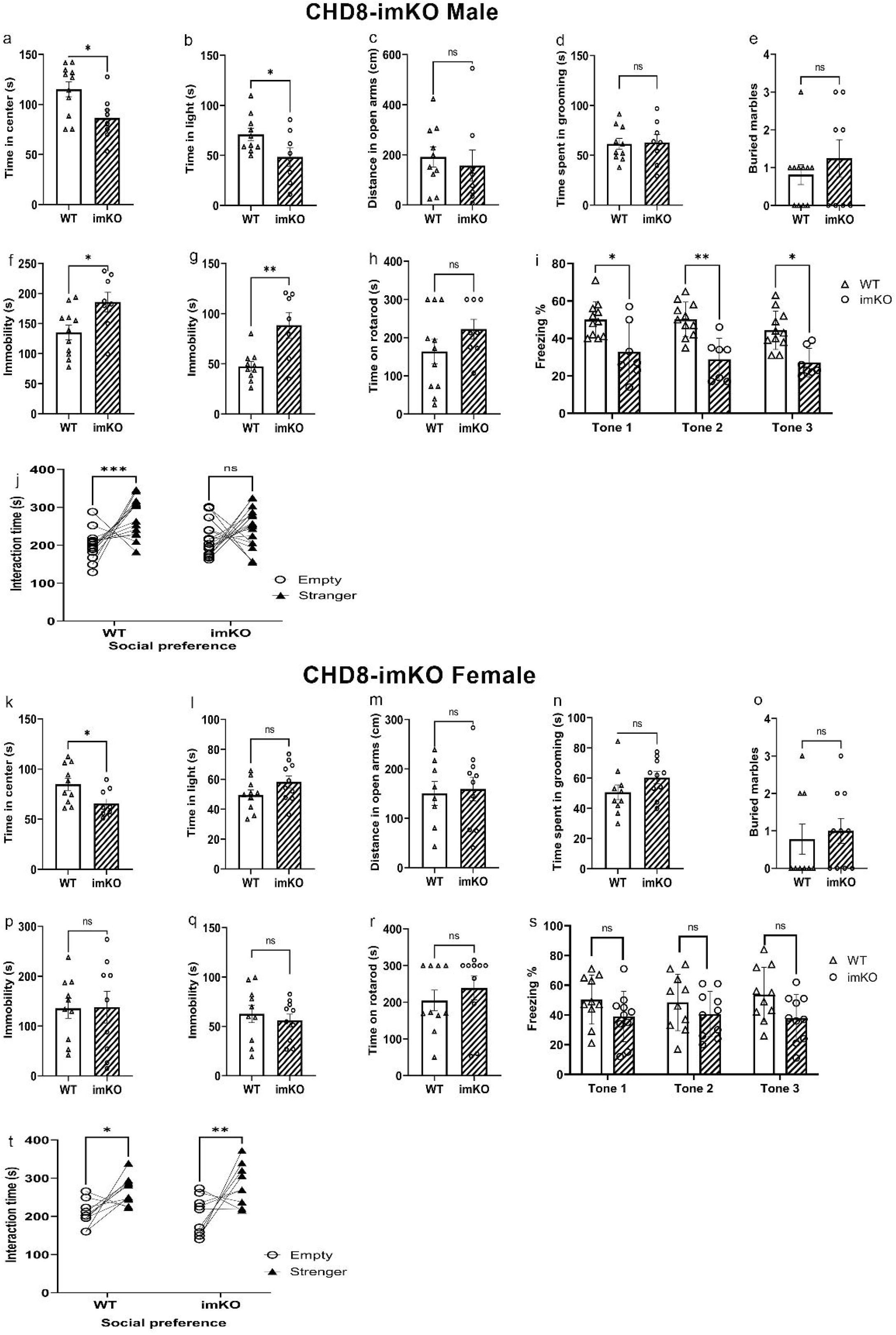
Behavioral Profiling of CHD8-imKO Mice. **CHD8-imKO males**: a) Time spent in the center of the open field. b) Time spent in the light area of the dark-light exploration test. c) Time spent in the open arm of the elevated plus maze. imKO male mice spent less time in the center of the open field and less time in the light chamber (*p=0.0205, *p=0.0473, respectively, two-way student’s t-test). d) Time self-grooming within a ten minute time period. e) Number of marbles buried in 30 minute time period. f) Time spent immobile in tail suspension test g) Time spent immobile in forced swim test. (*p=0.0195, **p=0.0037, two-way student’s t-test). h) Motor coordination and balance were assessed using the rotarod. Time spent before falling off rotarod. i) Percent freezing in each of three tones in cue fear conditioning test (2Way ANOVA, Tukey’s post hoc test, *p=0.0172, **p=0.0016, *p=0.0173) j) Time spent in interaction zone with stranger mice and empty chamber in three-chamber social interaction test (2Way ANOVA, Tukey’s post hoc test, ***p= 0.0001, ^ns^p=0.1589). **CHD8-imKO females**: k) Time spent in the center of the open field (*p=0.0239). l) Time spent in the light area of the dark-light exploration test. m) Time spent in the open arm of the elevated plus maze. n) Time self-grooming within a ten minute time period. o) Number of marbles buried in 30 minute time period. p) Time spent immobile in tail suspension test q) Time spent immobile in forced swim test. r) Motor coordination and balance were assessed using the rotarod. Time spent before falling off rotarod. s) Percent freezing in each of three tones in cue fear conditioning test t) Time spent in interaction zone with stranger mice and empty chamber in three-chamber social interaction test (2Way ANOVA *p=0.0169, **p=0.0027). All tests were normal (l-dark-light, m-elevated plus maze, p-tail suspension test, q-forced swimming test).

The same behavioral tests were performed in female imKO mice. Female imKO mice displayed decreased time spent in the center of the open field (Fig. 2k). However, in all other behavioral assessments, including repetitive behavior tests, depression-related tests, fear conditioning, and sociability tests, there were no significant differences between female imKO mice and their wild type controls (Fig. 2l-t). Therefore, CHD8 deletion in microglia during adulthood induces a strong behavioral phenotype specifically in males.

To study the role of CHD8 within excitatory neurons, we performed the same behavioral tests in inKO mice. InKO male mice displayed reduced exploration in the center of the open field (Fig. 3a) but no differences between phenotypes were displayed in the dark light (Fig. 3b), elevated plus maze (Fig. 3c), and grooming tests Figure 3d). However, there was a decrease in the number of buried marbles (Fig. 3e). Moreover, inKO male mice exhibited increased immobility during the forced swimming test (Fig. 3f) and tail suspension test (Fig. 3g), suggesting depression-like behavior. Notably, motor function assessments remained unaltered (Fig. 3h). No significant differences were observed in the cue fear conditioning test (Fig. 3i) or sociability test (Fig. 3j). These results lead us to conclude that inKO male mice specifically display behaviors of increased depression-like phenotype.

**Figure 3.**
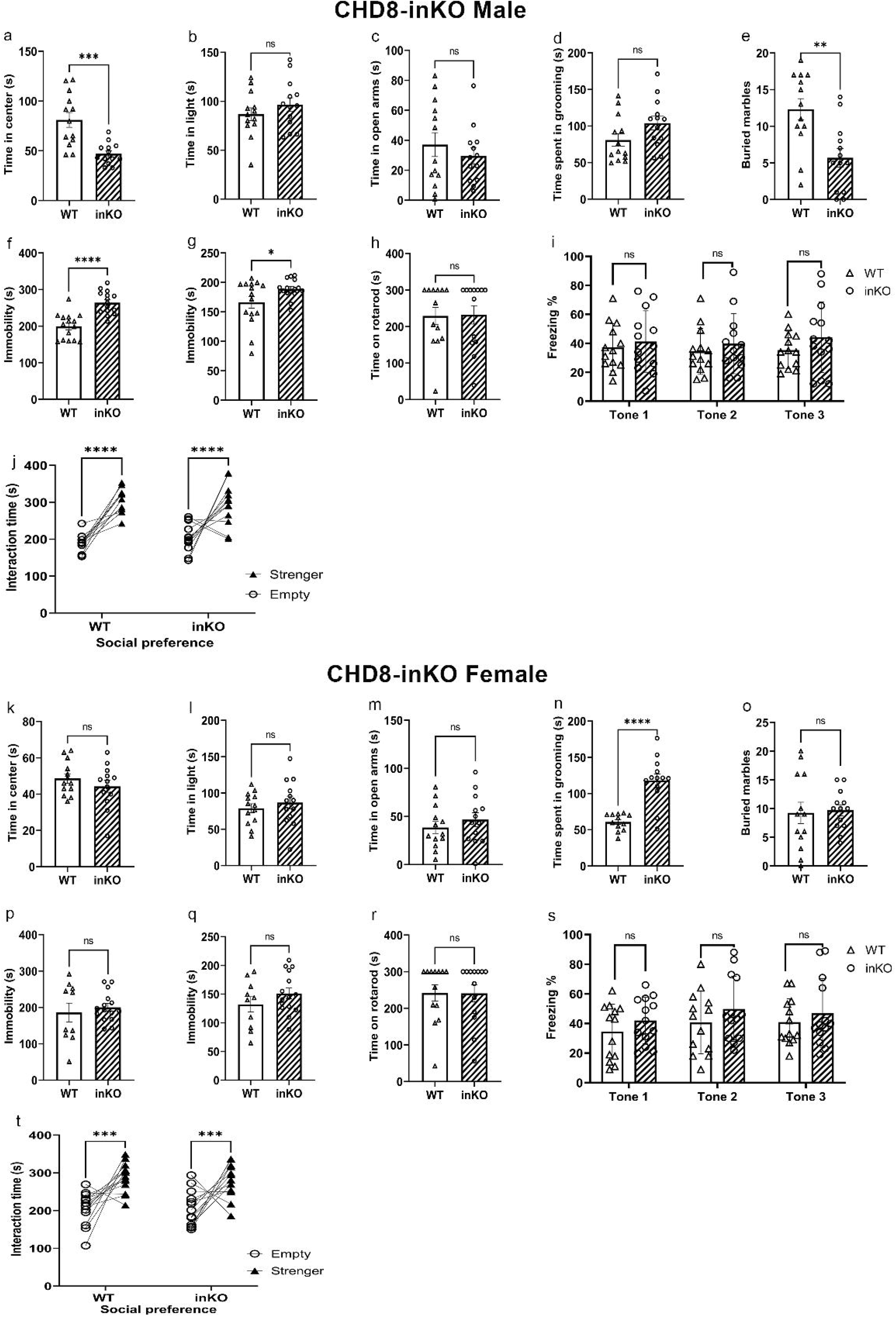
Behavioral Profiling of CHD8-inKO Mice. **CHD8-inKO male**: a) Time spent in the center of the open field. b) Time spent in the light area of the dark-light exploration test. c) Time spent in the open arm of the elevated plus maze. (***p=0.0003). d) Time self-grooming within a ten minute time period. e) Number of marbles buried in 30 minute time period. (**p=0.0022). f) Time spent immobile in tail suspension test (****p<0.0001) g) Time spent immobile in forced swim test (*p=0.0474). h) Motor coordination and balance were assessed using the rotarod. Time spent before falling off rotarod. i) Percent freezing in each of three tones in cue fear conditioning test j) Time spent in interaction zone with stranger mice and empty chamber in three-chamber social interaction test (2Way ANOVA ****p<0.0001, ****p<0.0001). **CHD8-inKO Female Mice**: k) Time spent in the center of the open field. l) Time spent in the light area of the dark-light exploration test. m) Time spent in the open arm of the elevated plus maze. n) Time self-grooming within a ten minute time period. (****p<0.0001). o) Number of marbles buried in 30 minute time period. p) Time spent immobile in tail suspension test q) Time spent immobile in forced swim test. r) Motor coordination and balance were assessed using the rotarod. Time spent before falling off rotarod. s) Percent freezing in each of three tones in cue fear conditioning test t) Time spent in interaction zone with stranger mice and empty chamber in three-chamber social interaction test (2Way ANOVA ***p=0.001, ***p=0.0006).

InKO female mice exhibited minor behavioral alterations characterized by a reduction in total distance moved during the open field test (Fig. S3a) and heightened grooming behaviors (Fig. 3n). Interestingly, assessments of other anxiety-related behaviors (Fig. 3k-m) and the number of buried marbles (Fig. 3o) in inKO female mice yielded results comparable to the wt group. In contrast to male mice, inKO female mice did not display any depression-like behaviors (Fig. 3p,q). Additionally, motor function assessments (Fig. 3r), cue fear conditioning tests (Fig. 3s), and sociability assessments (Fig. 3t) in female inKO mice were consistent with those of control groups. These results demonstrate a sex-specific imbalance in which CHD8 adulthood knockout in excitatory neurons induces depression-like behavior exclusively in male mice only.

### Sex-Specific Alterations in CHD8-imKO Mice: Microglial Morphology and Transcriptomic Profile

Microglia morphology was studied in male and female imKO mice. Male imKO mice exhibited a reduction in the number of microglial cells within the hippocampus and PFC (Fig. 4a,b,c). TUNEL staining determined no increase in apoptosis in male imKO mice, suggesting that the population reduction might not be attributed to apoptosis (Supplementary Figure 1). Moreover, microglial ramification was significantly decreased, evident as reduced intersection numbers from the soma (Fig. 4d) and lower dendritic length (Fig. 4e,j). A decrease in ramification is a hallmark of a pro-inflammatory state of the microglia. Conversely, imKO female mice did not display any significant differences in the microglial population (Fig. 4a,f,g) or characteristics (Fig. 4h,i,j). This analysis highlights the influence of CHD8 in microglia particularly in males.

**Figure 4.**
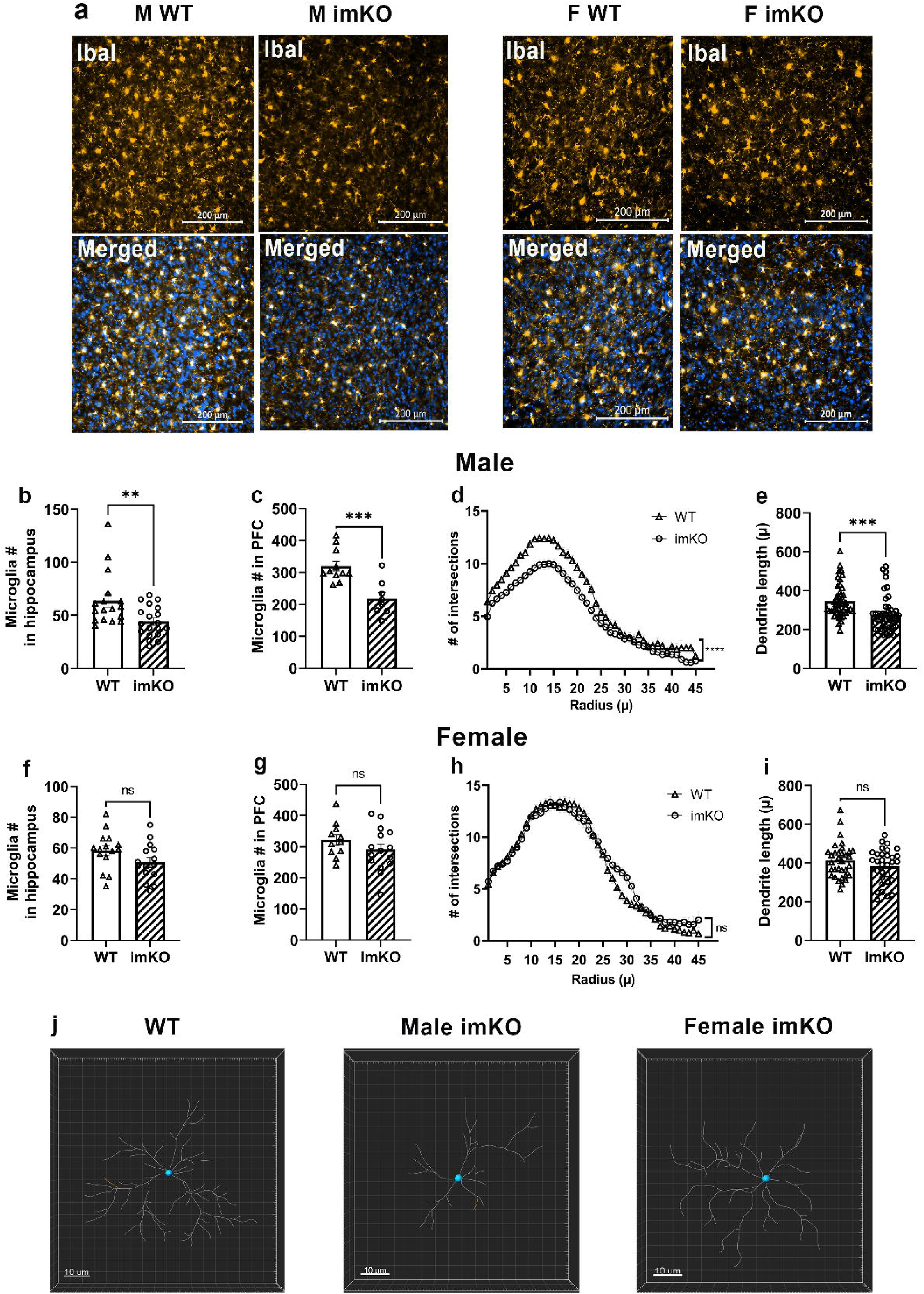
Microglial Morphology in CHD8-imKO Mice. Microglial morphology was assessed in the hippocampus and prefrontal cortex (PFC) of male and female CHD8-imKO mice. a) Representative pictures of IBA-1 positive cells in imKO and wt mice in male and female PFC. b) Microglia number (IBA-1+ cells) in hippocampus of male mice (**p=0.0070) c) IBA-1 positive cells in PFC of male mice (***p=0.0007). d) Sholl analysis and e) dendrite length of microglial cells in the PFC indicate a reduction in cell complexity (****p<0.0001 two-way ANOVA, ***p=0.0001). F) IBA-1+ cells in hippocampus of female mice. g) IBA-1+ cells in PFC of female mice h) Sholl analysis in the PFC of female mice i) Measurement of microglial dendrite length in the PFC of female mice. Representative Tracking of microglia dendrites in j) WT, k) male imKO, and l) female imKO mice using Imaris.

RNA sequencing was employed to study transcriptomic alterations in microglia after CHD8 depletion. Given the limited population of microglial cells in the brain, we performed FACS-mediated isolation of microglia from the entire brain, followed by whole transcriptome analysis (Fig. 5a). CD45+, TMEM119+, and Cd11b+ antibodies were used for identification and sorting of microglia (Fig. 5b). In male mice, there were a total of 1,381 genes that were differentially expressed between CHD8 imKO microglia and wild type microglia (Fig. 5c, Supplementary Table 1). In female mice, only 75 genes were differentially expressed between CHD8 imKO microglia and wild type microglia (Fig. 5d, Supplementary Table 2). Therefore, CHD8 had a more profound effect on gene expression in the microglia in males compared to females. The Gene Ontology (GO) analysis revealed many alterations in males, including the upregulation of chemokines, cytokines, cell cycle-related genes, ATPase activity-dependent DNA processes, and positive regulation of neuron development (Fig. 5e). Cytokine-related genes that were upregulated include *CCL2*, *CCL5*, C*CL12*, *CX3CL1*, among others. Cell cycle-related genes that were upregulated include multiple genes that downregulate cell cycle progression, including *CHEK*2 and *MTUS1*. These findings correlate with the increase in the inflammatory state of the microglia but a decrease in the number of activated microglia. Concurrently, there was a discernible downregulation of ribosomal and structural proteins, which could support the decrease in microglial numbers. In comparison, females displayed comparatively fewer pathway changes, mainly marked by the upregulation of ubiquitin and enzymatic binding pathways, alongside downregulation observed in ribosomal and translational genes (Fig. 5f). These findings underscore a distinct gender-specific response to CHD8 depletion in microglia, with males exhibiting a more substantial transcriptional reprogramming compared to females.

**Figure 5.**
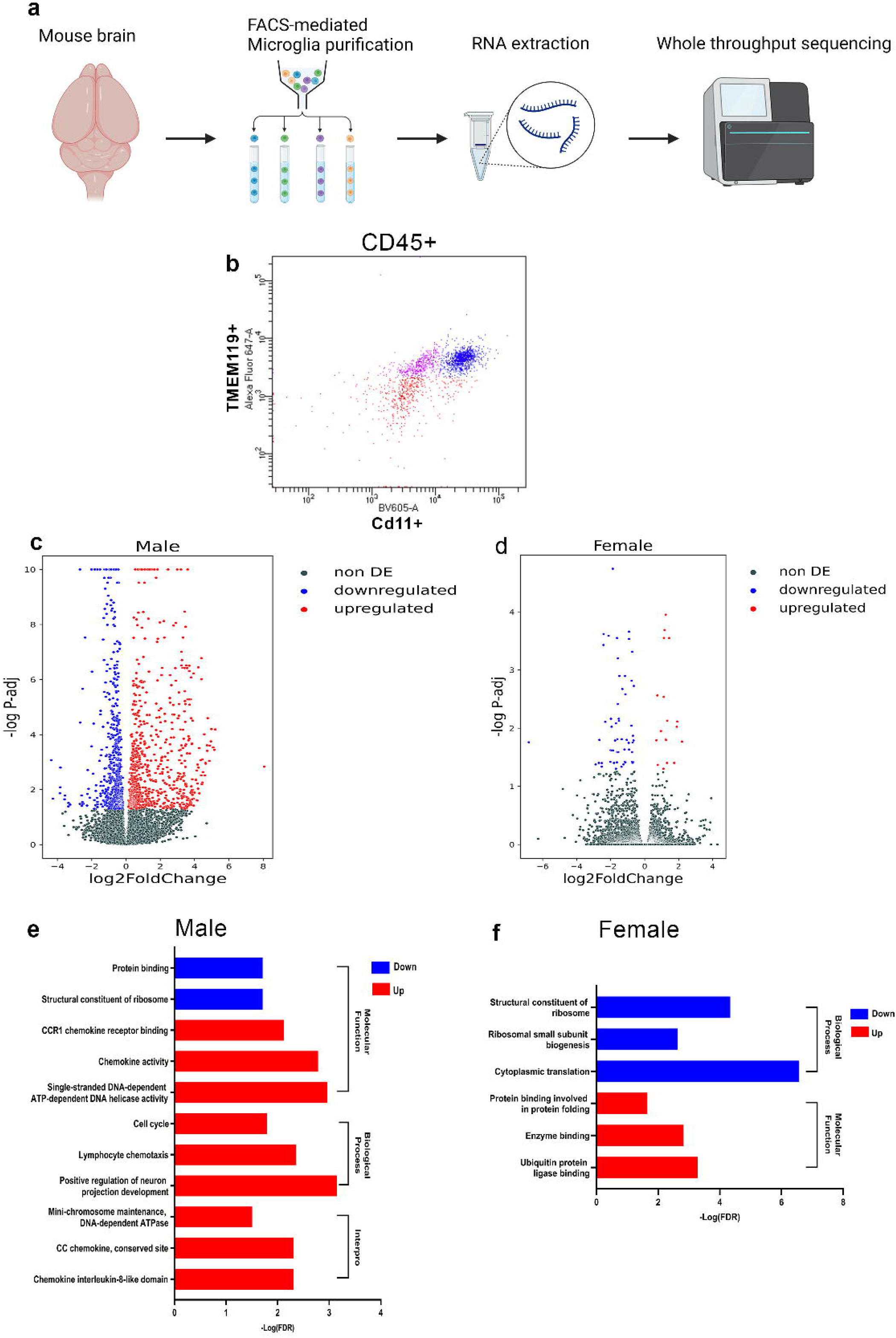
Differential Gene Expression Analysis in CHD8-imKO Mice. RNA-Seq analysis was performed on microglia from the whole brain tissues obtained from male and female CHD8-imKO mice. a) Microglia sorting from the whole brain and b) FACS analysis by three antibodies (CD45+, TMEM119+, Cd11b+) that together identify specifically microglia. In the graph, the presorted CD45+ population is presented, TMEM119 is represented on the Y axis, and CD11b on the X axis. Blue cells are TMEM+CD11b+, red cells are TMEM-CD11b-, and purple cells are TMEM+CD11b-. c,d) Volcano plot illustrating differentially expressed genes in the microglia of (c)male and (d)female CHD8-inKO mice. e,f) Gene ontology of differentially expressed genes in (e)male and (f)female imKO mice.

### Assessment of Neuronal Morphology and Transcriptomic Profiling in inKO Mice

While fewer behavioral changes were seen in the inKO mice, compared to the imKO mice, further experimentation was performed to understand the possible mechanisms underlying existing dysregulation in the inKO mice. GOLGI staining was used to analyze the neuronal morphology in both male and female inKO mice, evaluating dendrite length and complexity. Surprisingly, our findings revealed no significant alterations in neuron morphology in male or female inKO mice (Fig. 6). This analysis suggests that CHD8 depletion during adulthood does not significantly impact dendritic morphology.

**Figure 6.**
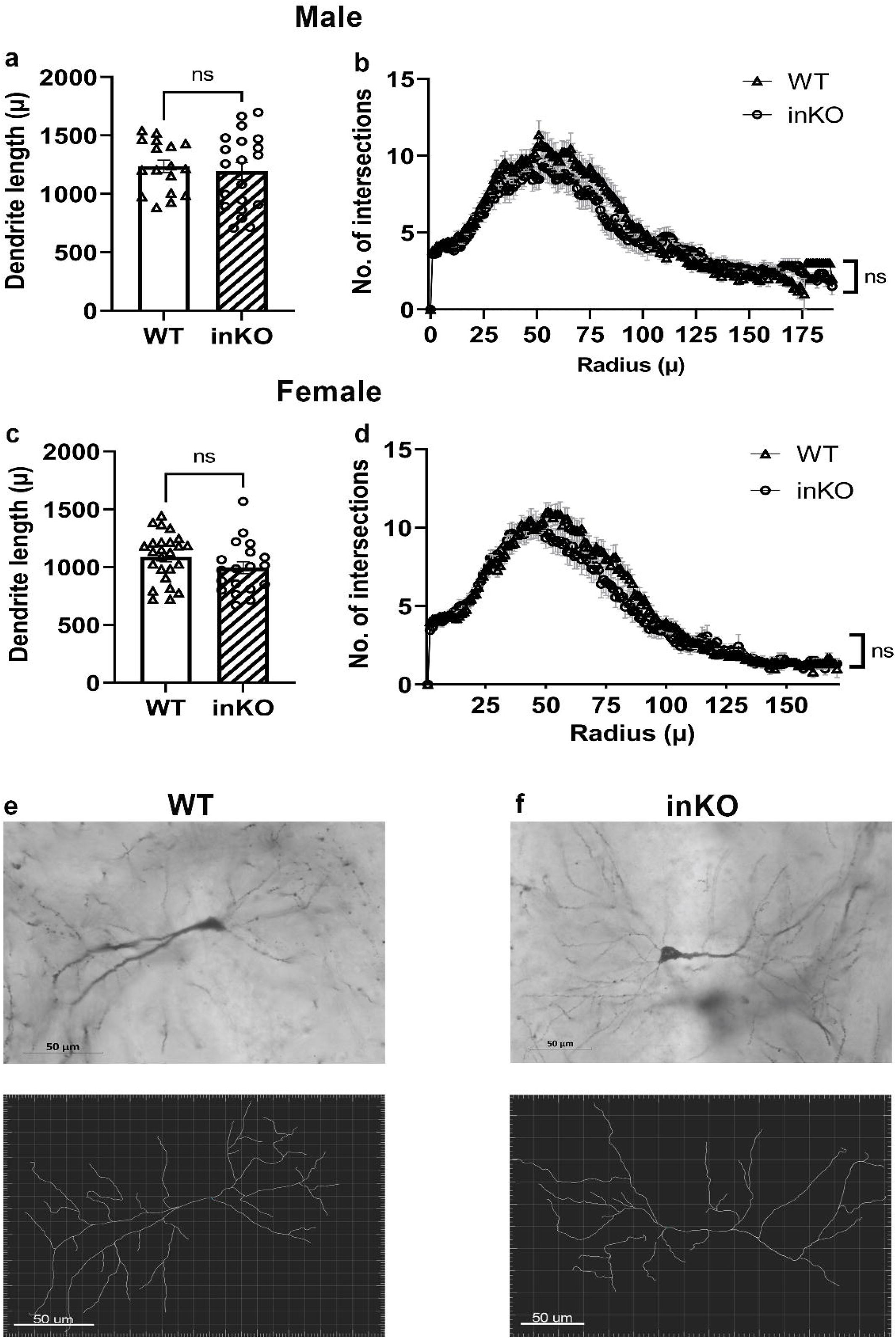
Morphological Analysis of Excitatory Neurons in CHD8-inKO Mice. a) Dendrite length in hippocampus in male wt and inKO mice b) Sholl analysis of dendritic complexity in male wt and inKO mice. c) Dendrite length in hippocampus in female wt inKO mice d) Sholl analysis of dendritic complexity in female wt and inKO mice Representative pictures and tracks of e) WT and f) inKO neurons.

The RNA-seq analysis of the hippocampus in inKO mice revealed only a few statistically significant changes between wt and inKO mice. There were only 10 differentially expressed genes in inKO males and six differentially expressed genes in inKO females (Fig. 7a,b, Supplementary Tables 3,4). These differentially expressed genes in both groups were not enriched in any gene ontology categories. Gene Set Enrichment Analysis (GSEA) was further performed on the dataset to discover if any biological pathways were enriched in genes with high fold change, regardless of the statistical significance of individual genes. GSEA discovered the heightened expression of genes associated with the Hedgehog and Wnt/Beta-catenin pathways specifically in male inKO mice (Fig. 7c,d). Remarkably, these alterations were absent in the hippocampal transcriptome of inKO female mice. These findings from RNA-Seq further strengthen sex-specific alterations in gene expression profiles in the hippocampus of CHD8-inKO mice. CHD8 in excitatory neurons affects behavior and gene expression during adulthood in males only.

**Figure 7.**
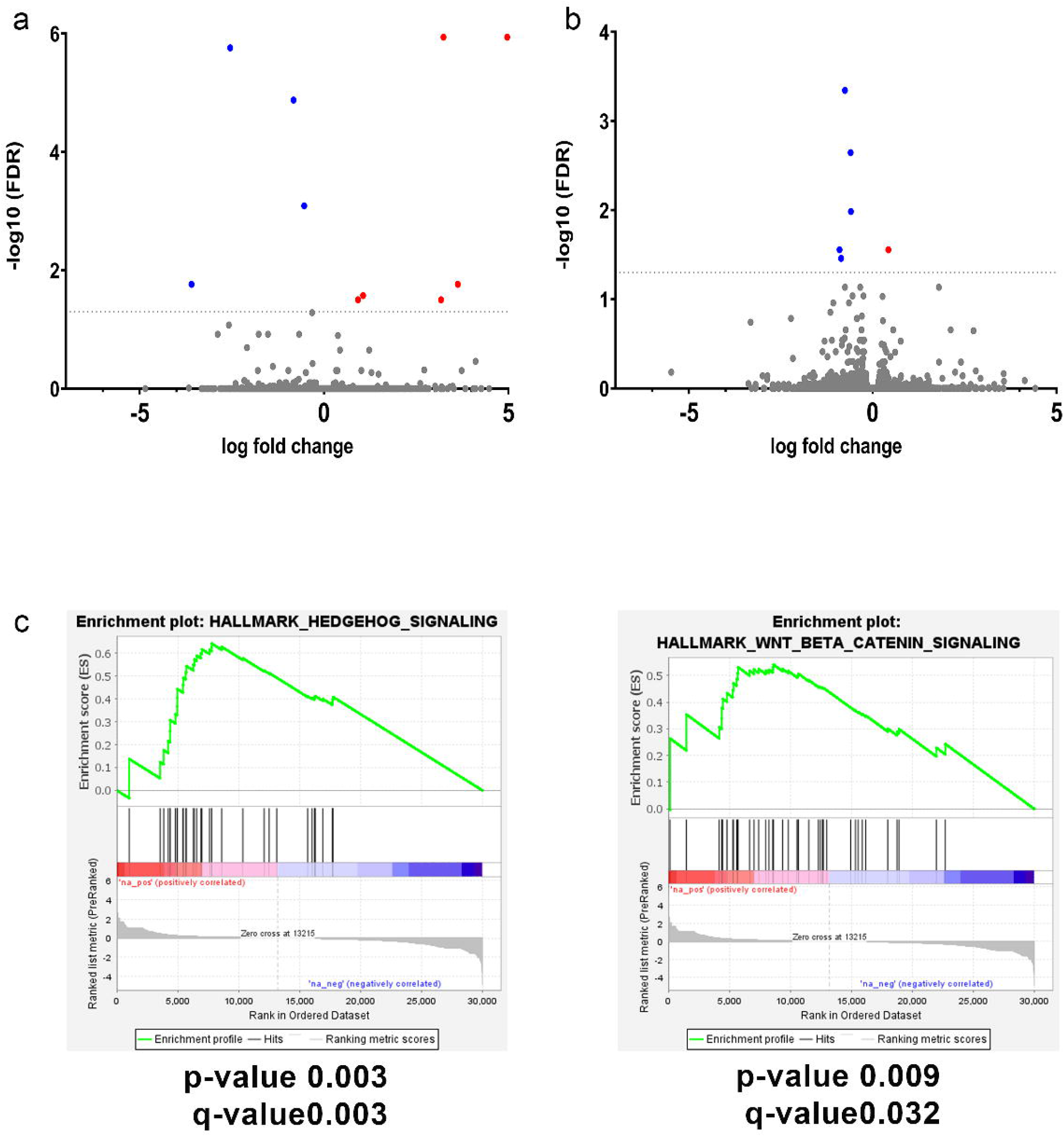
Differential Gene Expression Analysis in CHD8-inKO Mice. RNA-Seq analysis of hippocampal samples from CHD8-inKO male and female mice. Volcano plots highlight minor transcriptional alterations in CHD8-inKO (a) males and (b) females. (c) Gene Set Enrichment Analysis (GSEA) detects enrichment for genes within the Hedgehog and Wnt/Beta-catenin pathways that are upregulated specifically in males.

## Discussion

This study explored the sex-dependent role of CHD8 during adulthood in microglia and excitatory neurons. To achieve this, we generated conditional knockouts specifically in microglia and excitatory neurons during adulthood. We utilized CHD8^flox^/Cx3cr1^Cre-ERT+^ (imKO) mice for microglia and CHD8^flox^/CamKIIa^Cre-ERT+^ (inKO) mice for excitatory neurons. These newly developed mouse lines allowed us to study the involvement of CHD8 within distinct cell populations during adulthood, considering both male and female subjects.

Deletion of CHD8 had a more pronounced effect in males compared to females in both inKO and imKO mice. This was particularly apparent in the microglia, where male imKO displayed significantly higher levels of dysregulation compared to female imKO mice at the behavioral, cellular, and gene expression levels. Microglial depletion of CHD8 in males manifests in a comprehensive behavioral phenotype, characterized by impaired sociability, anxiety, deficient learning and memory, and depressive behaviors. Conversely, in inKO mice, CHD8 depletion in males primarily leads only to a depressive phenotype. Jung et al. previously observed that CHD8 depletion in Chd8+/N237k mice predominantly impacted males at the levels of behavior, neuronal function, and gene expression^28^.

Only a limited number of studies have explored CHD8’s influence on depression. Cherepanov et al. reported a depressive phenotype exclusively in female CHD8+/ΔSL mice, however, they used a haploinsufficiency developmental model ^29^. Most other studies have not included or reported depression-related tests in their behavioral tests. Therefore, we cannot rule out the possibility that these phenotypes might also be found in developmental models. Moreover, emerging evidence suggests a role for activated microglia in depression ^57^. In our gene expression analysis, we found an increase in several cytokines, including CCL2, in imKO mice. CCL2 has been implicated as a driver of depression-like behavior ^58^.

The current study reveals that CHD8 depletion in microglia induces both morphological and transcriptional alterations. Our results align with existing studies that have reported gene expression changes linked to immune response activation in the context of autism ^50,59^. Activated microglia exhibit a dynamic morphological transition between pro-inflammatory and anti-inflammatory stages, in which activated microglia display less branching, otherwise known as decreased ramification. Microglia characterized by a short ramification morphology release pro-inflammatory cytokines and chemokines, potentially contributing to neuronal damage ^46,49,60–62^.

CHD8 imKO microglia displayed decreased ramification at the morphological level, in parallel to increased transcription of cytokines. Therefore, CHD8 knockout induced an activation of microglia. In a separate study of CHD8 knockdown in human cerebral organoids, Astorkia et al. demonstrated that CDH8 reduction leads to impaired neuronal-glial communication ^63^. The decrease in number of microglia, in comparison to the activation of microglia, is more difficult to interpret. The transcriptional data determined dysregulation of several cell-cycle related genes, including genes responsible for inhibiting cell-cycle progression. This may be the reason for the fall in microglia number, particularly since there was no apoptosis detected in the tissue.

Multiple previous studies have found evidence for sex-specific differences in microglial function. Villa et al. studied microglia sex deferences in adult mice and found multiple gene expression alterations indicating the neuroprotective phenotype of microglia in females, compared to males. In addition, transplantation of female microglia in males helped to treat damage caused by acute cerebral ischemia^61^. Another study found that females have increased anti-inflammatory microglia during adulthood^62^. In addition, Xu et al. revealed that exaggerated protein translation in microglia impaired sociability behavior in males, not females, accompanied by microglia morphological changes and a reduction in the number of microglia in PFC in males only. RNA-sequencing found that this was accompanied by upregulation of cytokines and chemokines expression in microglia KO males only (Xu *et al.*, 2020). Therefore, male microglia seem to be particularly vulnerable to dysregulation, although it is still not completely clear why. A reason for the sex-specific effects of microglial CHD8 knockout would be the differential binding of CHD8 on genomic regions between male and female microglia. Future experimentation should look into possible sex-specific binding of CHD8 in microglia.

This study unveils gender-specific variations in both the morphology and transcriptional profile of microglia, offering pivotal insights into the consequences of CHD8 deficiency on microglial populations. In contrast to the negligible impact observed in excitatory neuron morphology and transcriptomic following CHD8 depletion, microglial depletion resulted in a reduction in cell number in the prefrontal cortex (PFC) and hippocampus. Notably, reduced ramification was observed exclusively in males, underscoring the sex-specific nature of these morphological changes. Moreover, a substantial alteration in gene expression was evident, with the effects being more pronounced in males.

Various CHD8 mouse models have reported changes in neuronal gene expression^28,31,34–36^, although the changes in adult animals are often very minimal. The current study revealed elevated gene transcription associated with Hedgehog and Wnt/β atenin signaling pathways according to the GSEA analysis. Wnt/β-catenin signaling is crucial for diverse processes in the central nervous system, including neuronal differentiation, synaptic maintenance, and axonal remodeling, has been associated with autism ^64^. CHD8 is a β-catenin binding protein, and several studies have shown that CHD8 is a negative regulator of the β-catenin pathway^25,65^. Some previous studies have also found that deletion of CHD8 is associated with upregulated inWnt/β-catenin signaling in neurons. Platt. et. al. also reported an increase in Wnt/β-catenin-associated gene transcription specifically in the nucleus accumbens of mice with a CHD8 mutation ^34^. While CHD8 is expressed at high levels in neurons during the embryonic stage, peaking at E16-18, its expression dramatically decreases postnatally ^34,68,69^. Along with the fact that neurons do not proliferate during adulthood, this may explain the limited impact of CHD8 depletion in adulthood on neuronal morphology and gene transcription.

This novel study determined how CHD8 affects behavior and gene transcription during adulthood and comparatively in microglia and neurons in a sex-specific manner. Interestingly, CHD8 depletion in microglia causes strong autism-like behaviors in adult males. On the other hand, CHD8 depletion in excitatory neurons mainly leads to a depression-like phenotype. Also, a male bias, as seen in humans, was observed in these mice models. This research gives us new information about how CHD8 can influence autism-related processes in adults, and its effect on specific cell populations.

## Methods

### Mouse Models

Mice were housed according to Federation of Laboratory Animal Science Associations (FELASA) guidelines. Mice were maintained in a vivarium at 22 C with a reversed light-cycle (lights on at 19:00 hours, off at 07:00 hours). Food and water were provided ad libitum, except during behavioral testing. Testing occurred between 9:00 and 16:00 hours. Littermate mice were randomly selected to experimental groups. All experimentation was approved by the Institutional Animal Care and Use Committee (IACUC) under protocol number 46-07-2021.

#### inKO Mouse Model

To induce conditional knockout of CHD8 in excitatory neurons, the CamKiia-Cre-Ert mouse line (kindly provided by the laboratory of Prof. Alon Chen, Weizmann Institute of Science, Israel) was crossed with CHD8^flx/flx^ mice (The Jackson Laboratory, Strain #:031555) to generate the CHD8^flx/flx^Camk2aCreErt^+/-^ cKO mice. Then, these were further bred with CHD8^flx/flx^ mice to produce both the CHD8^flx/flx^Camk2aCreErt^+/-^ (cKO) and CHD8^flx/flx^Camk2aCreErt^-/-^ (wild-type) mice, forming the basis for the experiments. All offspring maintained the CHD8^flx/flx^ genotype, with half carrying the Camk2aCreErt^+/-^ allele. The Camk2aCreErt^-/-^ mice served as control animals throughout the experiments.

#### imKO Mouse Model

In parallel, the Cx3cr1-Cre-Ert mouse line (kindly provided by the laboratory of Prof. Stephen Jung, Weizmann Institute of Science, ISrael) was used to specifically target CHD8 in microglia. These mice were crossed with CHD8^flx/flx^ mice (The Jackson Laboratory, Strain #:031555) to generate the CHD8^flx/flx^Cx3cr1CreErt^+/-^ (cKO) and CHD8^flx/flx^Cx3cr1CreErt^-/-^ (wild-type) mice. All experimental studies involved crossing these two lines, ensuring that all offspring maintained the CHD8^flx/flx^ genotype, with half carrying the Cx3cr1CreErt^+/-^ allele.

### DNA extraction and genotyping

Genotyping was carried out using DNA isolated from ear tissue followed by PCR. **DNA Extraction:** Ear tissue samples from mice were suspended for 30 minutes at 95°C in the DNA extraction buffer. The extraction buffer consisted of NaOH (5M) and EDTA (0.5M) at pH 7.6-8. Specific primers were utilized for each target gene: **CHD8 Primers:** Forward Primer: 5’ TGG GGT GCT GGG AAC AGT A 3’. Reverse Primer: 5’ GCT CAC ACG AAT ATA ACC TCA CA 3’. **Camk2a-Cre-Ert Primers:** Forward Primer: 5’ GGT TCT CCG TTT GCA CTC AGG A 3’. Reverse Primers: Reverse 1: 5’ CTG CAT GCA CGG GAC AGC TCT 3’. Reverse 2: 5’ GCT TGC AGG TAC AGG AGG TAG T 3’. **Cx3cr1-Cre-Ert Primers:** Common Forward Primer: 5’ ACG CCC AGA CTA ATG GTG AC 3’. Mutant Reverse Primers: 5’ GTT AAT GAC CTG CAG CCA AG 3’. PCR amplification was performed using Master Mix 2xPCRBIO HS Taq Mix Red (PBo45618-032-17).

### Tamoxifen Treatment for Inducing Cre-ERT Expression

To activate the Cre-ERT system, all groups were given Tamoxifen treatment at 8 weeks old including both cKO and wt (wild-type) mice. Tamoxifen was administered once daily for four consecutive days at a dosage of 100 mg/kg per day via oral gavage. Four weeks post-treatment immunofluorescence was carried out followed by behavioral tests.

#### Animal Brain Fixation

Animals were sacrificed and brains were fixed through perfusion. A 100μl injection of Petal veterinary solution (pentobarbitone sodium 200 mg/ml CTS) diluted with saline (1:10) was administered to the mice. Blood was washed out with PBS injection into the heart, followed by fixation using 4% Paraformaldehyde (PFA) injection. The fixed brains were extracted and incubated overnight in 4% PFA, followed by another overnight incubation in 30% sucrose with 4% PFA. Brain slices of 30μ thickness were obtained using a sliding microtome (HM430) and stored in PBS with sodium azide at 4°C.

#### Immunostaining Protocol

PBS washes were performed on the brain slices, followed by a 10-minute incubation in Sodium citrate buffer (10mM Sodium citrate, 0.05% Tween 20, pH 6.0) for antigen retrieval. Further washing steps included PBST (0.3% Triton) and PBS. Blocking was carried out using normal horse serum for 1 hour at room temperature. Overnight incubation at 4°C was performed with primary antibodies diluted to 1:200 in PBS. After three washes with PBST and PBS, slices were incubated with secondary antibodies for 1 hour. Subsequent washes were performed, and the slices were mounted on glass slides using immu-mount. Throughout the procedure, incubations were conducted in shaking mode. Imaging was conducted using an Upright Microscope ApoTome or Slide Scanner.

Neurons were labeled using the anti-NeuN Antibody (clone A60, MAB377 Sigma-Aldrich, Host-mouse) as the primary antibody and Cy3 (Donkey anti-mouse IgG, Jackson, 715165150) as the secondary antibody. Microglia was marked by anti-IbaI (Goat anti-mouse IgG, Abcom, ab5076) as the primary antibody and Donkey anti-goat Cy3 as a secondary antibody. For CHD8, the anti-CHD8 antibody (ab84527, Rabbit polyclonal to CHD8, Isotype: IgG) served as the primary antibody, while Alexa Fluor® 488 (Goat anti-rabbit, Jackson, 11545144) was used as the secondary antibody. Nuclei were marked with Hoechst stain (Sigma, 1:10,000).

#### Behavioral tests

Mice, aged between 8-10 weeks, were administered a 100 mg/kg tamoxifen solution per day via gavage. Four weeks post the initial tamoxifen administration, behavioral tests were carried out. A camera was employed to record the mice’s movements. Tracking and analysis of animal behavior were conducted using EthoVision XT 10-Noldus.

#### Open Field Test and Grooming

The open field test was performed to evaluate anxiety-like behavior in mice within a novel environment. Each mouse was placed in a 50×50×30cm open field box, positioned in a corner, under a light intensity of approximately ∼120 lux. The first 10 minutes of the test were analyzed for cumulative duration spent in the center and border zones, along with the total distance moved by the mouse. The subsequent 10 minutes of the session were dedicated to manually assessing grooming behaviors.

#### Dark-Light Test

The dark-light test was conducted as an additional measure to evaluate anxiety-related behaviors in mice. A two-chambered cage setup was utilized for this test, consisting of a dark chamber made from black plastic with a closable top, and a light chamber made from white plastic that remained open at the top. The light intensity in the light chamber was maintained at approximately ∼1200 lux. Initially, the mouse was placed in the dark chamber with the top cover closed. Access to the light chamber was provided by opening the entryway, allowing the mouse to freely move between chambers for 10 minutes. The analysis involved tracking the frequency and cumulative duration of the mouse’s presence within the light arena. The latency to the first entry into the light chamber was also recorded as a measure of exploratory behavior.

#### Elevated Plus Maze Test

This test was used to evaluate anxiety-related behaviors in mice, focusing on their exploration of the safe and unsafe zones of the maze. An X-shaped apparatus comprising two open arms (30cm by 5 cm, white plastic) and two closed arms (15 cm high walls) was utilized for the test. The maze was positioned 55cm above the floor, maintaining a light intensity of 20 lux. The mouse was initially placed in the central square (5 by 5 cm) of the maze and allowed to navigate among the arms for 5 minutes. Analysis included the frequency and cumulative duration of visits to the center, open, and closed arms. Additionally, parameters such as latency to the first arm entry and the total distance moved during the test were recorded.

### Marble Burying Test

The marble burying test was carried out to evaluate repetitive behaviors exhibited by mice. The cage was lined with a 5 cm layer of sawdust, and 20 marbles of uniform size and color were evenly arranged on the surface. Lighting conditions were maintained at 20 lux throughout the test. The mouse was placed in a corner of the cage and allowed to explore the environment freely for 30 minutes. Marbles that were buried to a depth ranging between 70 to 100% beneath the sawdust layer were counted.

#### Tail Suspension Test

During the tail suspension test, the mouse was suspended by the last cm of its tail, allowing it to hang at 60 cm above the floor within a chamber (20□×□40□×□60 cm). Video recording captured the mouse behavior during the 6-minute test, focusing on total immobility.

#### Forced Swimming Test

A dark chamber was filled with water at room temperature to a depth of 15 cm. The mouse was placed in the water for a total duration of 6 minutes, with the first 2 minutes dedicated to habituation, and was video recorded. The last 4 minutes of the test were analyzed, focusing on the observation of mouse immobility.

#### Rotarod Test

Mice were placed on a rotating cylinder apparatus (Med Associates, St. Albins, VT) set at a speed of 40 rpm, with a maximum test duration of 300 seconds. The time that each mouse remained on the rotating cylinder was measured as a marker of motor function.

#### Wire Hanging Test

Mice were given a 2 mm thick horizontal metal wire to grip onto. The test was performed five times for each mouse to ensure accuracy and consistency. The time until the mouse fell from the wire, was recorded during each trial. Based on the five trials, an average latency to fall was calculated to represent its grip motor functions.

#### Beam Test-6 and 12 mm Thickness

Mice were placed on a 1-meter-long horizontal beam, varying in thickness between 6 mm and 12 mm. The time taken by the mouse to reach a designated safety zone from the starting point on the beam was measured. Additionally, the number of falls the mouse experienced while traversing the beam was recorded. Before the test, mice underwent a training phase where they were familiarized with navigating through the beam. Training sessions were carried out over two consecutive days, with mice undergoing two sessions per day to ensure familiarity and consistency.

#### Cue Fear Conditioning Test

The cue fear conditioning test was utilized to study memory and learning abilities in mice. The initial day was allocated for training and habituation. Mice were introduced into the test chamber (10.5 × 10.5 × 10.5 cm) for five minutes. A 75 dB tone served as the conditioned stimulus for 30 seconds, followed by a brief, mild foot shock (0.7 mA) lasting two seconds as the unconditioned stimulus. After a one-minute interval, another tone-shock pair was administered, and subsequently, the mouse was returned to its home cage one minute after the second tone-shock pairing. The following day, mice were placed in a different chamber with novel sensory cues (odor, flooring, and lighting) for cue-dependent memory assessment. After a two-minute habituation period, the tone was presented three times for 30 seconds each, with one-minute intervals between tones. Freezing behavior during the three-tone presentations was recorded as a measure of cue-dependent memory recall.

### Sociability Assessment

The three-chambered social test aimed to evaluate sociability behavior in mice through social interaction assessment. The testing apparatus consisted of a three-chambered cage divided by two entrances, each chamber measuring 20 by 40 by 22 cm. A clear plastic cylinder was positioned between the right and left chambers to facilitate controlled social interaction. Two mice of identical genotypes were introduced into the clear plastic cylinder for 10 minutes each day over 5 days for habituation purposes. The subject mouse was positioned in the center of the middle chamber. Initially, an unfamiliar mouse (stranger) was placed in the right cylinder for a 5-minute habituation phase. Following this habituation, gates providing access to the right and left chambers were opened, enabling the tested mouse to freely explore all three chambers for 10 minutes. Behavioral metrics included measuring the time and distance spent by the tested mouse in each chamber and within a predefined sniffing zone.

### Golgi Staining for Excitatory Neuron Morphology

GOLGI-Cox stain kit from Bioenno Tech, LLC was employed to analyze the morphology of excitatory neurons following the manufacturer’s protocol. 12-week-old mice brains were extracted and freshly dissected brains were sectioned at 180 µm thickness using a vibratome (Leica VT 1200s). Excitatory neurons were randomly selected for analysis, and their images were captured using a bright-field microscope. IMARIS 9.1.2 software was utilized to measure dendritic length and conduct Sholl analysis for neuron morphology assessment. Statistical analysis was performed using two-way ANOVA via GraphPad Prism software.

### Sample Collection and Neuronal RNA Sequencing

Brains from 12-week-old mice were harvested and washed with PBS buffer and the hippocampal tissue was dissected. RNA extraction was performed utilizing the RNeasy Micro Kit (QIAGEN, 74004) following the manufacturer’s protocol. A total of 30 libraries, accounting for 20.84 Gbp, were generated and subjected to initial quality control analysis. Quality assessment was conducted, and the samples were trimmed accordingly. A secondary quality evaluation of the reads was performed. Fastq-Screen software (v0.14.1) was utilized to evaluate contamination by aligning randomly selected reads to various reference genomes. Reads were mapped to the mouse genome (GCF_000001635.27_GRCm39 assembly) using the STAR software (v.27.9a). Uniquely mapped reads were quantified based on their corresponding transcripts using FeatureCounts software (V2.0.3). Assessment of DNA contamination was conducted by analyzing mapped reads based on their genomic location (Exons/Introns). Removal of PCR duplicates was executed using the DupRadar r-package to ensure data integrity. Raw counts obtained were employed to identify differentially expressed genes across the experimental groups (FcKO, FWT, McKO, and MWT). Differential expression analysis was performed using the EdgeR R-package (v3.23.5) with genes filtered based on LogFC and FDR criteria (LogFC ±1, Adjusted p-value < 0.05). GSEA software was used for gene network analysis. Genes were preranked by logfold2 change and then subject to GSEA analysis.

### TUNEL Staining

The In-Situ Cell Death Detection Kit (Roche Life Science) was used to identify apoptotic processes within the brain tissue. 30 µm thick floating sections obtained from 4% formaldehyde-fixed brains were utilized for the staining analysis. The staining procedure followed the instructions provided by the manufacturer.

### Microglial Sorting via FACS

A 12-week-old mouse was anesthetized, and brains were extracted following PBS perfusion. Brain tissues underwent enzymatic digestion by incubating at 37°C with 1ml of digestion buffer for 20 minutes (pipet after 10 min), followed by passage through a grey mesh to homogenize. After enzymatic digestion, the sample was washed using 10ml MACS buffer. Gentlely pounding of the mesh with a syringe’s rubber end assisted tissue passage. To halt collagenase action, a total of 20-30ml MACS buffer was added, followed by centrifugation at 2200rpm, 4°C for 5 minutes. The pellet was resuspended in 1ml of 40% percol solution and transferred to a tube containing 2ml of 40% percol, totaling 3ml in a 15ml tube. Centrifugation was performed at 900g, room temperature, no breaks for 15 minutes. Percol was aspirated, resuspended the pellet in ∼100ul MACS buffer, avoiding debris, and transferred through a white mesh into a fresh FACS tube containing 5ml cold MACS buffer on ice. Subsequent centrifugation was conducted at 1400rpm, 4°C for 5 minutes. The supernatant was discarded, and the pellet was gently vortexed. Fc blocks (10 ul of 1:20 in FACS buffer) were added and incubated for 15 minutes on ice. Extracellular staining was performed without washing by adding 50 ul of antibodies (CD45-Alexa 488, CD11b-BV 605, Tmem119-Alexa 647 at 1:100 dilution in MACS buffer) and incubating for 15 minutes on ice in the dark. Washed cells were centrifuged at 1400rpm, 4°C for 5 minutes. The supernatant was discarded, and the pellet was resuspended in 300ul MACS buffer. The cell suspension was transferred into a new FACS tube and sorted using Aria-Sorter based on specific antibodies (CD45, CD11b, Tmem119) identifying microglia.

### Microglial RNA Sequencing

Following cell sorting, RNA was extracted from the isolated microglial cells. RNA sequencing was performed using the MARS-seq2.0 methodology, following the protocol described by Keren-Shaul et al ^56^.

---

**Supplementary Figure 1**

**TUNEL staining.**

TUNEL analysis revealed no evidence of apoptosis in the PFC, as demonstrated by nuclei staining in blue (Hoechst) and the absence of DNA fragments in green (fluorescein).

## Acknowledgments

We thank the Israel National Center for Personalized Medicine in the Weizmann Institute of Science for help with microglial RNA sequencing. This study was funded by grants from the Israel Science Foundation (898/17 and 1159/22).

## References

1. Pina-Camacho, L. et al. Autism spectrum disorder: does neuroimaging support the DSM-5 proposal for a symptom dyad? A systematic review of functional magnetic resonance imaging and diffusion tensor imaging studies. J. Autism Dev. Disord. 42, 1326–1341 (2012).

2. Kereszturi, É. Diversity and classification of genetic variations in autism spectrum disorder. Int. J. Mol. Sci. 24, (2023).

3. Maenner, M. J. et al. Prevalence and Characteristics of Autism Spectrum Disorder Among Children Aged 8 Years - Autism and Developmental Disabilities Monitoring Network, 11 Sites, United States, 2018. MMWR Surveill. Summ. 70, 1–16 (2021).

4. Choueiri, R., Garrison, W. T. & Tokatli, V. Early identification of autism spectrum disorder (ASD): strategies for use in local communities. Indian J. Pediatr. 90, 377–386 (2023).

5. Botsas, G., Koidou, E., Chatzinikolaou, K. & Grouios, G. Environmental Influences on Individuals with Autistic Spectrum Disorders with Special Emphasis on Seasonality: An Overview. Children (Basel) 10, (2023).

6. Höfer, J. et al. Pathways to a diagnosis of autism spectrum disorder in Germany: a survey of parents. Child Adolesc. Psychiatry Ment. Health 13, 16 (2019).

7. Howsmon, D. P., Kruger, U., Melnyk, S., James, S. J. & Hahn, J. Classification and adaptive behavior prediction of children with autism spectrum disorder based upon multivariate data analysis of markers of oxidative stress and DNA methylation. PLoS Comput. Biol. 13, e1005385 (2017).

8. Wang, L., Wang, B., Wu, C., Wang, J. & Sun, M. Autism spectrum disorder: neurodevelopmental risk factors, biological mechanism, and precision therapy. Int. J. Mol. Sci. 24, (2023).

9. Wing, L. & Gould, J. Severe impairments of social interaction and associated abnormalities in children: epidemiology and classification. J. Autism Dev. Disord. 9, 11–29 (1979).

10. Weissberg, O. & Elliott, E. The mechanisms of CHD8 in neurodevelopment and autism spectrum disorders. Genes 12, (2021).

11. Thompson, B. A., Tremblay, V., Lin, G. & Bochar, D. A. CHD8 is an ATP-dependent chromatin remodeling factor that regulates beta-catenin target genes. Mol. Cell. Biol. 28, 3894–3904 (2008).

12. Zahir, F. et al. Novel deletions of 14q11.2 associated with developmental delay, cognitive impairment and similar minor anomalies in three children. J. Med. Genet. 44, 556–561 (2007).

13. Neale, B. M. et al. Patterns and rates of exonic de novo mutations in autism spectrum disorders. Nature 485, 242–245 (2012).

14. Wilkinson, B. et al. The autism-associated gene chromodomain helicase DNA-binding protein 8 (CHD8) regulates noncoding RNAs and autism-related genes. Transl. Psychiatry 5, e568 (2015).

15. Satterstrom, F. K. et al. Large-Scale Exome Sequencing Study Implicates Both Developmental and Functional Changes in the Neurobiology of Autism. Cell 180, 568–584.e23 (2020).

16. Lee, S. Y., Kweon, H., Kang, H. & Kim, E. Age-differential sexual dimorphism in CHD8-S62X-mutant mouse behaviors. Front. Mol. Neurosci. 15, 1022306 (2022).

17. Kweon, H. et al. Excitatory neuronal CHD8 in the regulation of neocortical development and sensory-motor behaviors. Cell Rep. 34, 108780 (2021).

18. Bernier, R. et al. Disruptive CHD8 mutations define a subtype of autism early in development. Cell 158, 263–276 (2014).

19. O’Roak, B. J. et al. Sporadic autism exomes reveal a highly interconnected protein network of de novo mutations. Nature 485, 246–250 (2012).

20. O’Roak, B. J. et al. Multiplex targeted sequencing identifies recurrently mutated genes in autism spectrum disorders. Science 338, 1619–1622 (2012).

21. An, Y. et al. De novo variants in the Helicase-C domain of CHD8 are associated with severe phenotypes including autism, language disability and overgrowth. Hum. Genet. 139, 499–512 (2020).

22. Douzgou, S. et al. The clinical presentation caused by truncating CHD8 variants. Clin. Genet. 96, 72–84 (2019).

23. Wang, T. et al. De novo genic mutations among a Chinese autism spectrum disorder cohort. Nat. Commun. 7, 13316 (2016).

24. Stolerman, E. S., Smith, B., Chaubey, A. & Jones, J. R. CHD8 intragenic deletion associated with autism spectrum disorder. Eur. J. Med. Genet. 59, 189– 194 (2016).

25. Nishiyama, M., Skoultchi, A. I. & Nakayama, K. I. Histone H1 recruitment by CHD8 is essential for suppression of the Wnt-β-catenin signaling pathway. Mol. Cell. Biol. 32, 501–512 (2012).

26. Cotney, J. et al. The autism-associated chromatin modifier CHD8 regulates other autism risk genes during human neurodevelopment. Nat. Commun. 6, 6404 (2015).

27. Cerase, A. & Avner, P. From X-inactivation to neurodevelopment: CHD8-transcription factors (TFs) competitive binding at regulatory regions of CHD8 target genes can contribute to correct neuronal differentiation. Current Research in Neurobiology 5, 100114 (2023).

28. Jung, H. et al. Sexually dimorphic behavior, neuronal activity, and gene expression in Chd8-mutant mice. Nat. Neurosci. 21, 1218–1228 (2018).

29. Cherepanov, S. M. et al. Oxytocin ameliorates impaired social behavior in a Chd8 haploinsufficiency mouse model of autism. BMC Neurosci. 22, 32 (2021).

30. Stessman, H. A. F. et al. Targeted sequencing identifies 91 neurodevelopmental-disorder risk genes with autism and developmental-disability biases. Nat. Genet. 49, 515–526 (2017).

31. Jiménez, J. A. et al. Chd8 haploinsufficiency impairs early brain development and protein homeostasis later in life. Mol. Autism 11, 74 (2020).

32. Ostrowski, P. J. et al. The CHD8 overgrowth syndrome: A detailed evaluation of an emerging overgrowth phenotype in 27 patients. Am. J. Med. Genet. C Semin. Med. Genet. 181, 557–564 (2019).

33. Katayama, Y. et al. CHD8 haploinsufficiency results in autistic-like phenotypes in mice. Nature 537, 675–679 (2016).

34. Platt, R. J. et al. Chd8 Mutation Leads to Autistic-like Behaviors and Impaired Striatal Circuits. Cell Rep. 19, 335–350 (2017).

35. Gompers, A. L. et al. Germline Chd8 haploinsufficiency alters brain development in mouse. Nat. Neurosci. 20, 1062–1073 (2017).

36. Suetterlin, P. et al. Altered Neocortical Gene Expression, Brain Overgrowth and Functional Over-Connectivity in Chd8 Haploinsufficient Mice. Cereb. Cortex 28, 2192–2206 (2018).

37. Zhao, C. et al. Dual requirement of CHD8 for chromatin landscape establishment and histone methyltransferase recruitment to promote CNS myelination and repair. Dev. Cell 45, 753–768.e8 (2018).

38. Hulbert, S. W. et al. A novel chd8 mutant mouse displays altered ultrasonic vocalizations and enhanced motor coordination. Autism Res. 13, 1685–1697 (2020).

39. Kawamura, A. et al. Oligodendrocyte dysfunction due to Chd8 mutation gives rise to behavioral deficits in mice. Hum. Mol. Genet. 29, 1274–1291 (2020).

40. Hurley, S. et al. Distinct, dosage-sensitive requirements for the autism-associated factor CHD8 during cortical development. Mol. Autism 12, 16 (2021).

41. Kierdorf, K. & Prinz, M. Microglia in steady state. J. Clin. Invest. 127, 3201–3209 (2017).

42. Lukens, J. R. & Eyo, U. B. Microglia and neurodevelopmental disorders. Annu. Rev. Neurosci. 45, 425–445 (2022).

43. Fan, G., et al. Microglia modulate neurodevelopment in autism spectrum disorder and schizophrenia. Int. J. Mol. Sci. 24, (2023).

44. Schwarz, J. M., Sholar, P. W. & Bilbo, S. D. Sex differences in microglial colonization of the developing rat brain. J. Neurochem. 120, 948–963 (2012).

45. Carson, M. J., Doose, J. M., Melchior, B., Schmid, C. D. & Ploix, C. C. CNS immune privilege: hiding in plain sight. Immunol. Rev. 213, 48–65 (2006).

46. Yang, R. et al. Regulation of microglial process elongation, a featured characteristic of microglial plasticity. Pharmacol. Res. 139, 286–297 (2019).

47. McDougle, C. J. et al. Toward an immune-mediated subtype of autism spectrum disorder. Brain Res. 1617, 72–92 (2015).

48. Fernández de Cossío, L., Guzmán, A., van der Veldt, S. & Luheshi, G. N. Prenatal infection leads to ASD-like behavior and altered synaptic pruning in the mouse offspring. Brain Behav. Immun. 63, 88–98 (2017).

49. Morgan, J. T. et al. Microglial activation and increased microglial density observed in the dorsolateral prefrontal cortex in autism. Biol. Psychiatry 68, 368–376 (2010).

50. Gupta, S. et al. Transcriptome analysis reveals dysregulation of innate immune response genes and neuronal activity-dependent genes in autism. Nat. Commun. 5, 5748 (2014).

51. McCarthy, M. M. & Wright, C. L. Convergence of sex differences and the neuroimmune system in autism spectrum disorder. Biol. Psychiatry 81, 402–410 (2017).

52. Xiao, L. et al. Critical Role of TLR4 on the Microglia Activation Induced by Maternal LPS Exposure Leading to ASD-Like Behavior of Offspring. Front. Cell Dev. Biol. 9, 634837 (2021).

53. Zhao, D. et al. Transcriptome analysis of microglia in a mouse model of Rett syndrome: differential expression of genes associated with microglia/macrophage activation and cellular stress. Mol. Autism 8, 17 (2017).

54. Suzuki, K. et al. Microglial activation in young adults with autism spectrum disorder. JAMA Psychiatry 70, 49–58 (2013).

55. Schaafsma, W. et al. Maternal inflammation induces immune activation of fetal microglia and leads to disrupted microglia immune responses, behavior, and learning performance in adulthood. Neurobiol. Dis. 106, 291–300 (2017).

56. Keren-Shaul, H. et al. MARS-seq2.0: an experimental and analytical pipeline for indexed sorting combined with single-cell RNA sequencing. Nat. Protoc. 14, 1841–1862 (2019).

57. Huang, C. et al. The ketone body metabolite β droxybutyrate induces an antidepression-associated ramification of microglia via HDACs inhibition-triggered Akt-small RhoGTPase activation. Glia 66, 256–278 (2018).

58. Curzytek, K. & Leś iewicz, M. Targeting the CCL2-CCR2 axis in depressive disorders. Pharmacol. Rep. 73, 1052–1062 (2021).

59. Voineagu, I. et al. Transcriptomic analysis of autistic brain reveals convergent molecular pathology. Nature 474, 380–384 (2011).

60. Benedusi, V. et al. Liver ERα regulates AgRP neuronal activity in the arcuate nucleus of female mice. Sci. Rep. 7, 1194 (2017).

61. Villa, A. et al. Sex-Specific Features of Microglia from Adult Mice. Cell Rep. 23, 3501–3511 (2018).

62. McCarthy, M. M., Pickett, L. A., VanRyzin, J. W. & Kight, K. E. Surprising origins of sex differences in the brain. Horm. Behav. 76, 3–10 (2015).

63. Astorkia, M., Liu, Y., Pedrosa, E. M., Lachman, H. M. & Zheng, D. Molecular and network disruptions in neurodevelopment uncovered by single cell transcriptomics analysis of CHD8 heterozygous cerebral organoids. BioRxiv (2023) doi:10.1101/2023.09.27.559752.

64. Caracci, M. O. et al. Wnt/β-Catenin-Dependent Transcription in Autism Spectrum Disorders. Front. Mol. Neurosci. 14, 764756 (2021).

65. Kobayashi, M. et al. Nuclear localization of Duplin, a beta-catenin-binding protein, is essential for its inhibitory activity on the Wnt signaling pathway. J. Biol. Chem. 277, 5816–5822 (2002).

66. Wang, P. et al. CRISPR/Cas9-mediated heterozygous knockout of the autism gene CHD8 and characterization of its transcriptional networks in neurodevelopment. Mol. Autism 6, 55 (2015).

67. Wang, P. et al. CRISPR/Cas9-mediated heterozygous knockout of the autism gene CHD8 and characterization of its transcriptional networks in cerebral organoids derived from iPS cells. Mol. Autism 8, 11 (2017).

68. Xu, Q. et al. Autism-associated CHD8 deficiency impairs axon development and migration of cortical neurons. Mol. Autism 9, 65 (2018).

69. Durak, O. et al. Chd8 mediates cortical neurogenesis via transcriptional regulation of cell cycle and Wnt signaling. Nat. Neurosci. 19, 1477–1488 (2016).

