## Supplementary figures and images for "CHD8 adulthood microglial knockout induces behavioral, morphological, and transcriptional changes in a sex-dependent manner"

### Supplementary Figure 1

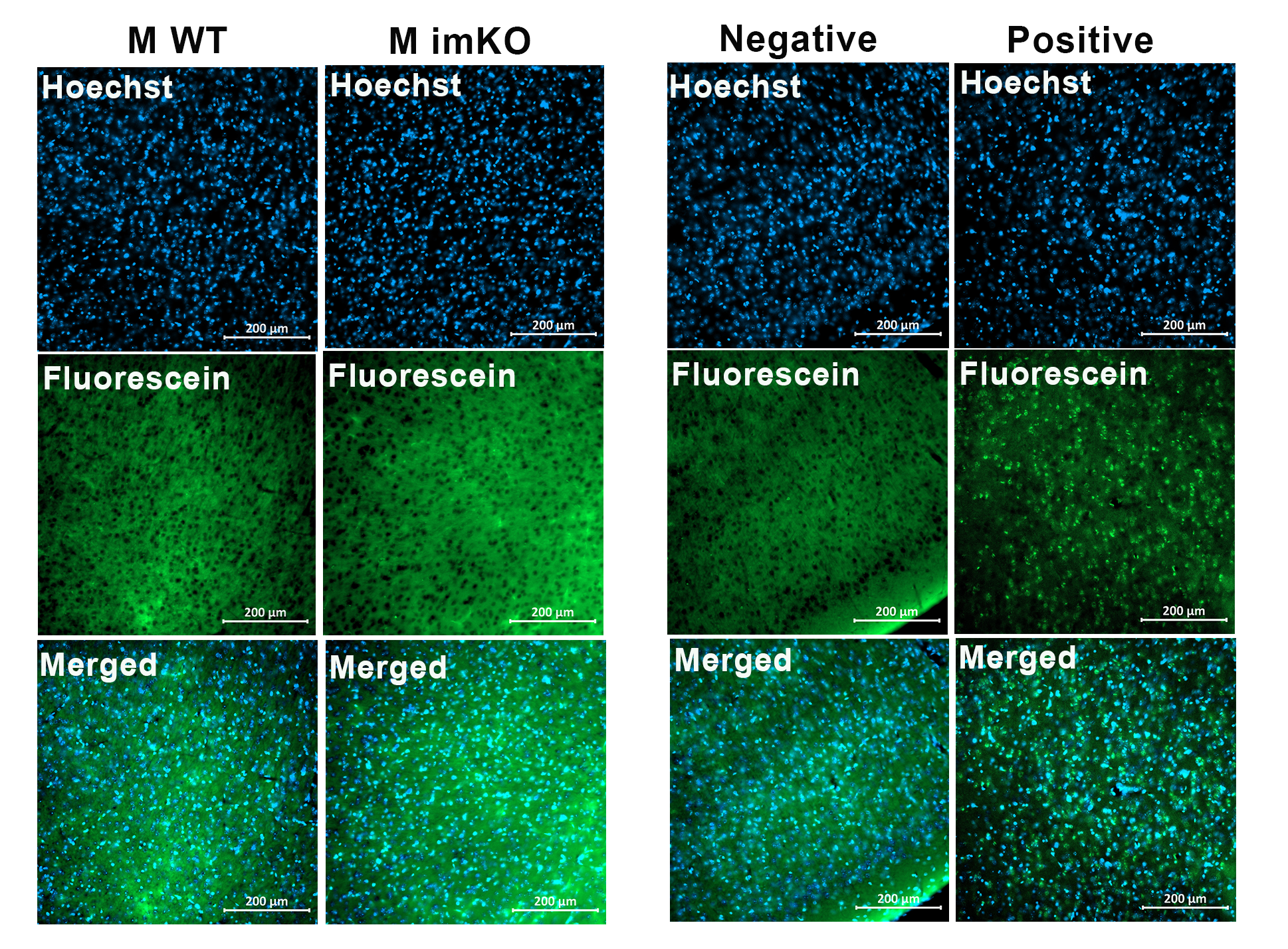
